# Comments on: “A comprehensive repertoire of tRNA-derived fragments in prostate cancer”

**DOI:** 10.1101/061572

**Authors:** Rogan Magee, Phillipe Loher, Aristeidis G. Telonis, Yohei Kirino, Isidore Rigoutsos

## Abstract

We evaluated the deep-sequencing (RNA-seq) data from human prostate tissue that were reported in [1] and the tRNA-derived fragments described in the original analysis. Our study of the same RNA-seq datasets reveals a considerably different pool of tRNA fragments, many of them with higher abundances than the fragments reported in [1]. We also evaluated the q-PCR approach proposed in [1]. As the approach lacks 5’-endpoint specificity, it will not estimate correctly the abundance of many of the tRFs that are present in the sampled RNA populations from human prostate tissue.

---

TRNA-derived fragments (tRFs) represent a new class of small non-coding RNAs that has been receiving increasing attention in recent years [2]. Several groups, including ours, have been reporting on the tRFs’ involvement in important processes such as inhibition of translation, RNA silencing, targeting of methylation, proliferation, promotion of metastasis, etc. [3–6]. A recent report by Olvedy et al [1] leveraged deep-sequencing data to generate and report a profile of tRNA fragments from 11 sources of human prostate tissue: formalin-fixed paraffin-embedded (FFPE) samples, and fresh frozen samples from normal adjacent tissue, benign prostatic hypertrophy group, and eight categories of prostate cancer (PCa).

## Many of the reported tRFs receive low or no support from the available RNA-seq data or map to non-tRNA sequences

For each of the 11 sources, RNA from 3-4 samples was extracted, pooled, and deep-sequenced. After adapter removal and quality-trimming, Olvedy et al mapped the sequenced reads while allowing mismatches on a composite database comprising the mature human cytosolic tRNAs from the gtRNAdb database, and the 22 human mitochondrial tRNAs, and reported 598 tRFs [1]. However, based on the data included in [1], 29 of the 598 reported tRFs received either zero support or support by a single read only; these are listed in Table 1, which is adapted from Supp. Table 3 of [1].

**Table 1.**
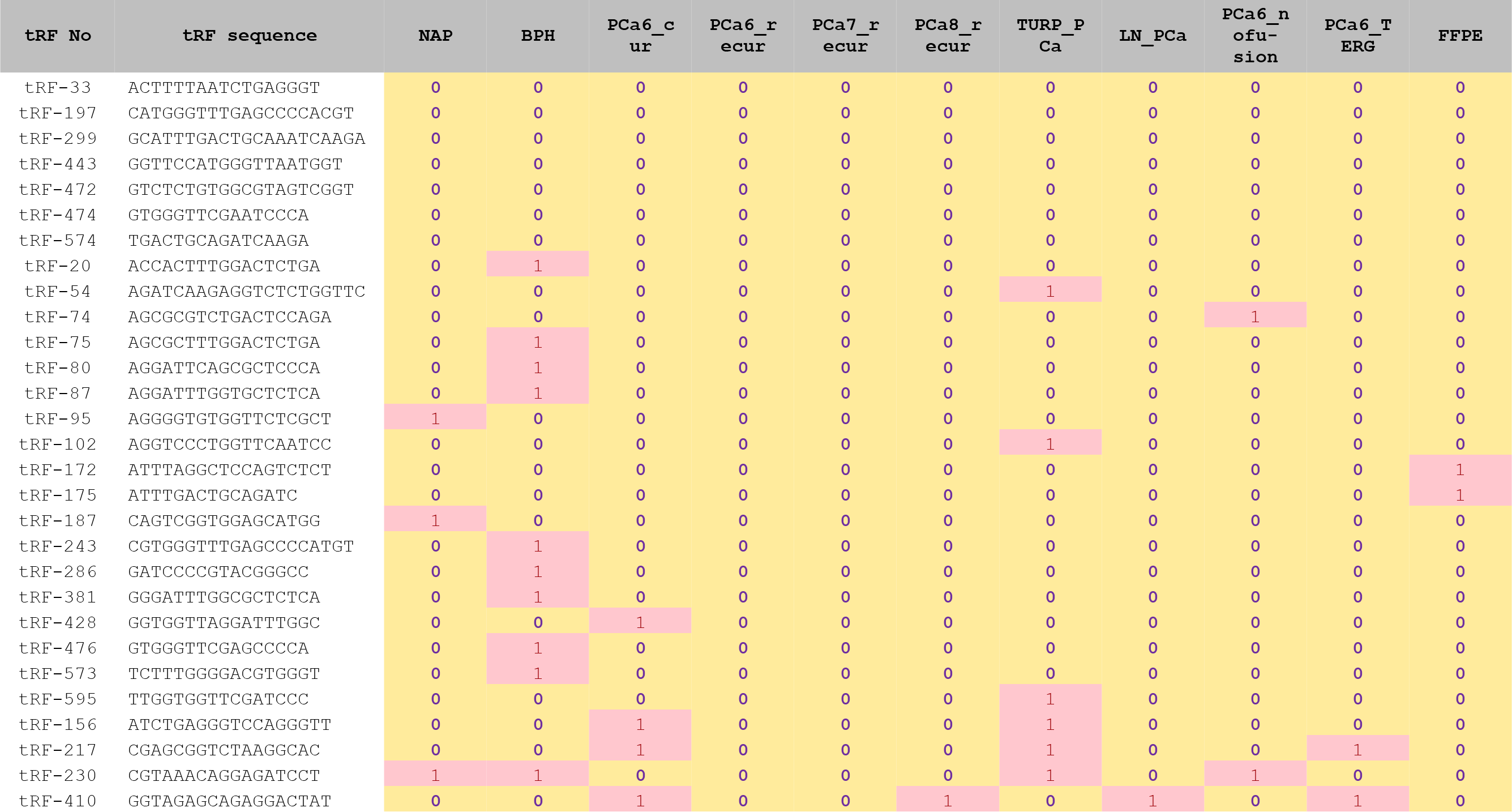
Some of the reported tRFs receive very low or zero support. The below table comprises rows sub-selected from Supp. Table 3 of [1] and shows that 29 of the 598 reported tRFs received very low or zero support in each of the 11 profiled datasets. Counts are in raw reads. The first column lists the tRF labels as they are used in [1].

In [5, 7] we discussed how exact mapping, i.e. mapping without indels or mismatches, assists in better tackling the cross-talk that results from the fact that near-similar nucleotide segments are present in multiple isodecoders of the same anticodon, in different isodecoders, or even in non-tRNA sequences. Using the method that we described in [5] we mapped the RNA-seq data of [1] anew and found that 192 (32.1%) of the originally reported 598 tRFs do not receive any support when indels or mismatches are not permitted during mapping (Table 2).

**Table 2.**
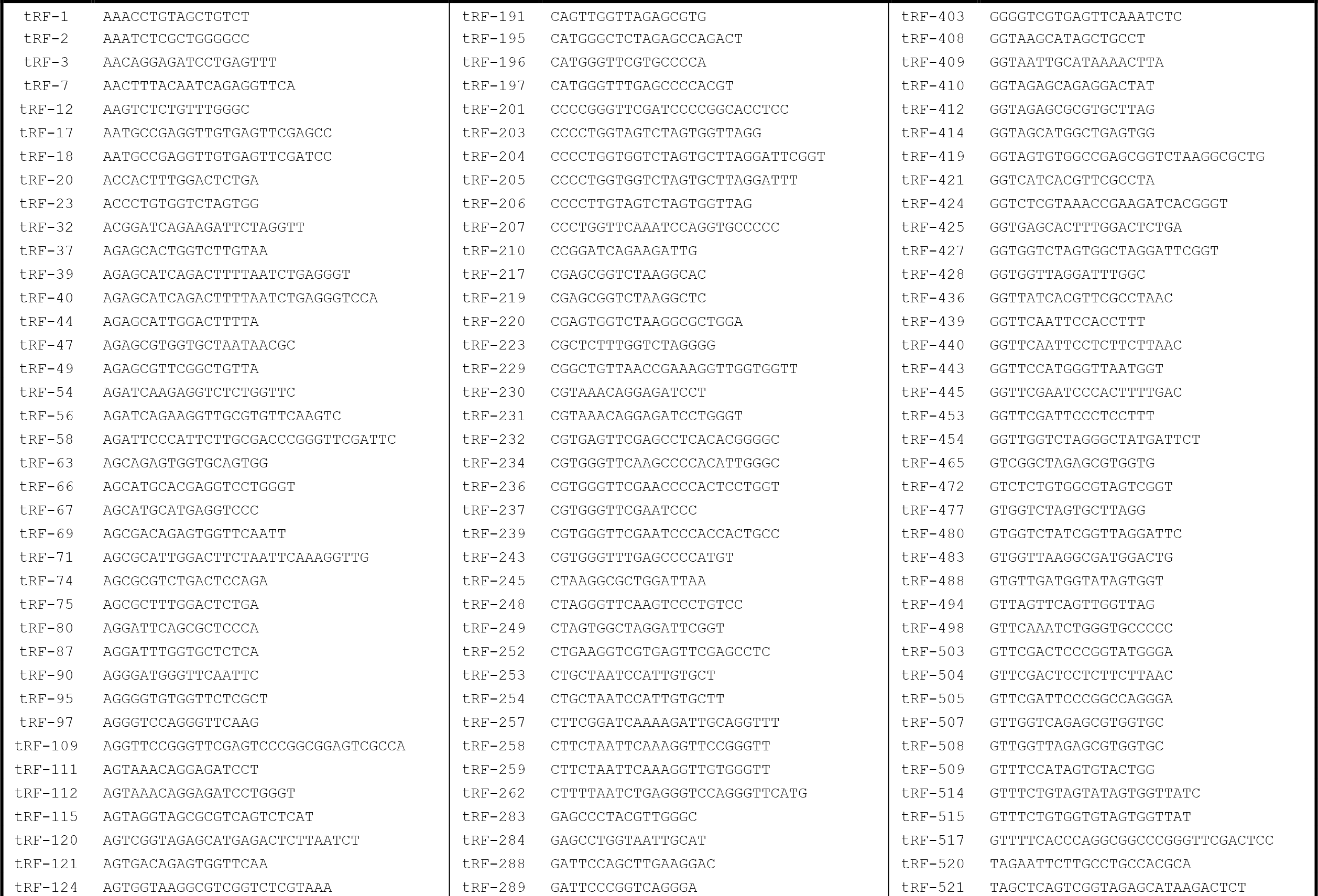
tRFs that do not receive support under exact mapping. A list of 192 tRFs from [1] with length of 16 nucleotides or longer that receive no support when exact mapping (no indels, no replacements) is used. Preceding each tRF is its label (e.g. tRF-1, tRF-191, tRF-403, etc.) as introduced in [1].

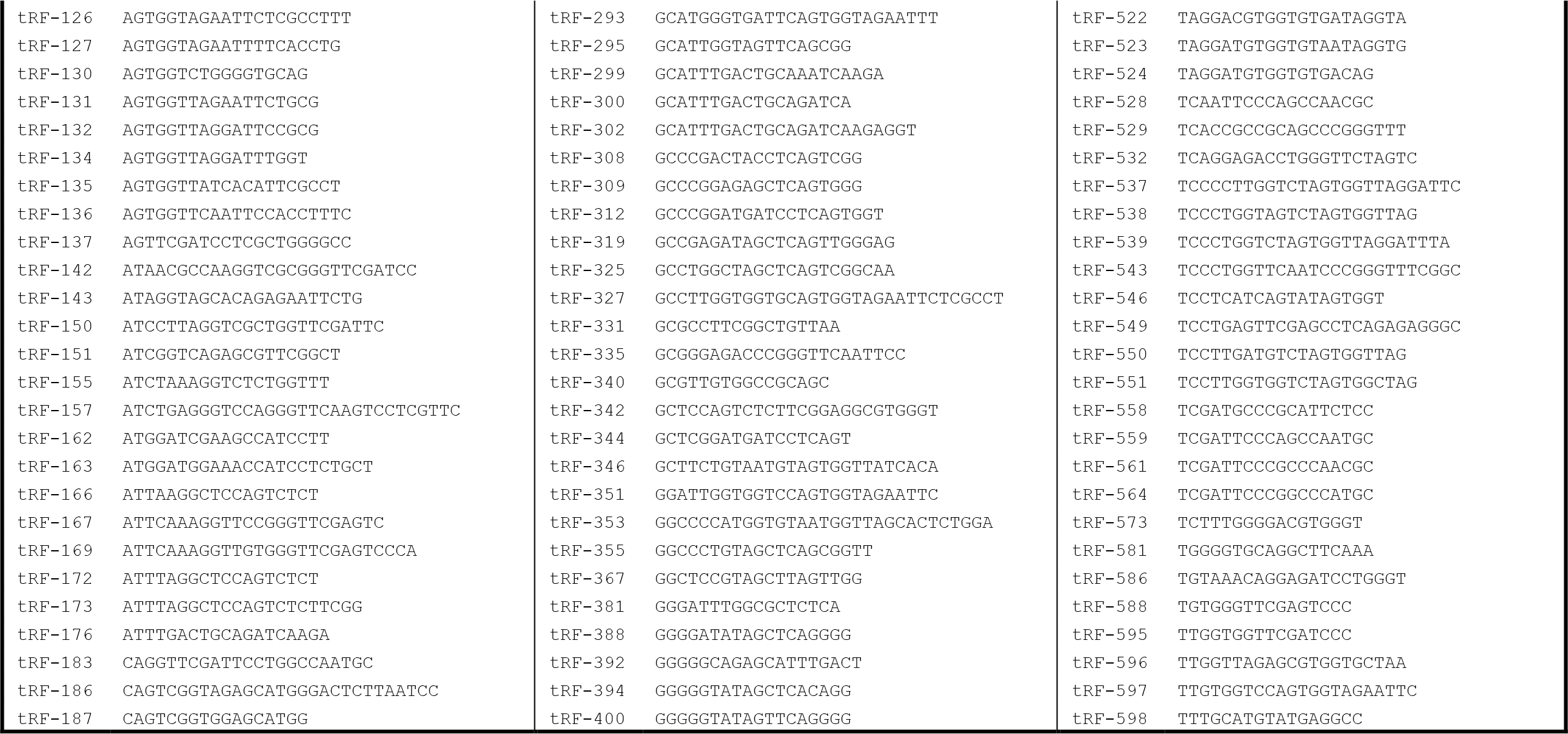

To avoid reporting as tRFs molecules that have non-tRNA origins, it is also important that reads be mapped to the *entire* human genome [5, 7], and not to the tRNA space alone as was done in [1]. In Supp. Table 1, we list 63 tRFs among the 598 reported in [1] whose exact copies can also be found at loci that are neither tRNAs nor any of the hundreds of partial tRNAs permeating the human genome [5, 7] and include repeat element sequences L1, L2, ERV1, ERVL, Alu, hAT-Tip100, etc. Among these 63 tRFs, tRF-125, tRF-171, tRF-298, and tRF-376 have characteristically high numbers of exact copies inside these non-tRNA categories of repeats.

## The generated RNA-seq datasets support numerous abundant tRFs that were not reported

We used a thresholding scheme that was adaptive and stringent, keeping from each dataset only the 3% most-abundant short RNAs among those reported by the method of [5] that carries out exact mapping on the full genome. The tRNA space is the same as the one used by Olvedy et al in [1]. At this stringent threshold setting, the RNA-seq data of [1] support 3,341 tRFs of which only 183 are among the originally reported collection of 598 tRFs. Curiously, most of the remaining 3,158 tRFs (94.5%) that were not reported in [1] are more abundant than the ones that were reported (Supp. Table 2) and present in at least two or more of the 11 datasets (Supp. Table 3). Of the 3,341 tRFs, 2,207 map exclusively to tRNA space whereas 1,134 map both inside and outside tRNA space (Supp. Table 2). Table 3 below lists the top-20 most abundant tRFs (in terms of the number of reads that support them) for each of the 11 RNA-seq datasets; the complete list appears in Supp. Table 2. Table 4 shows the 195 of the 3,341 tRFs that appear in all 11 of the analyzed datasets: of these, only 37 were reported in [1] – see also Supp. Table 3.

**Table 3.**
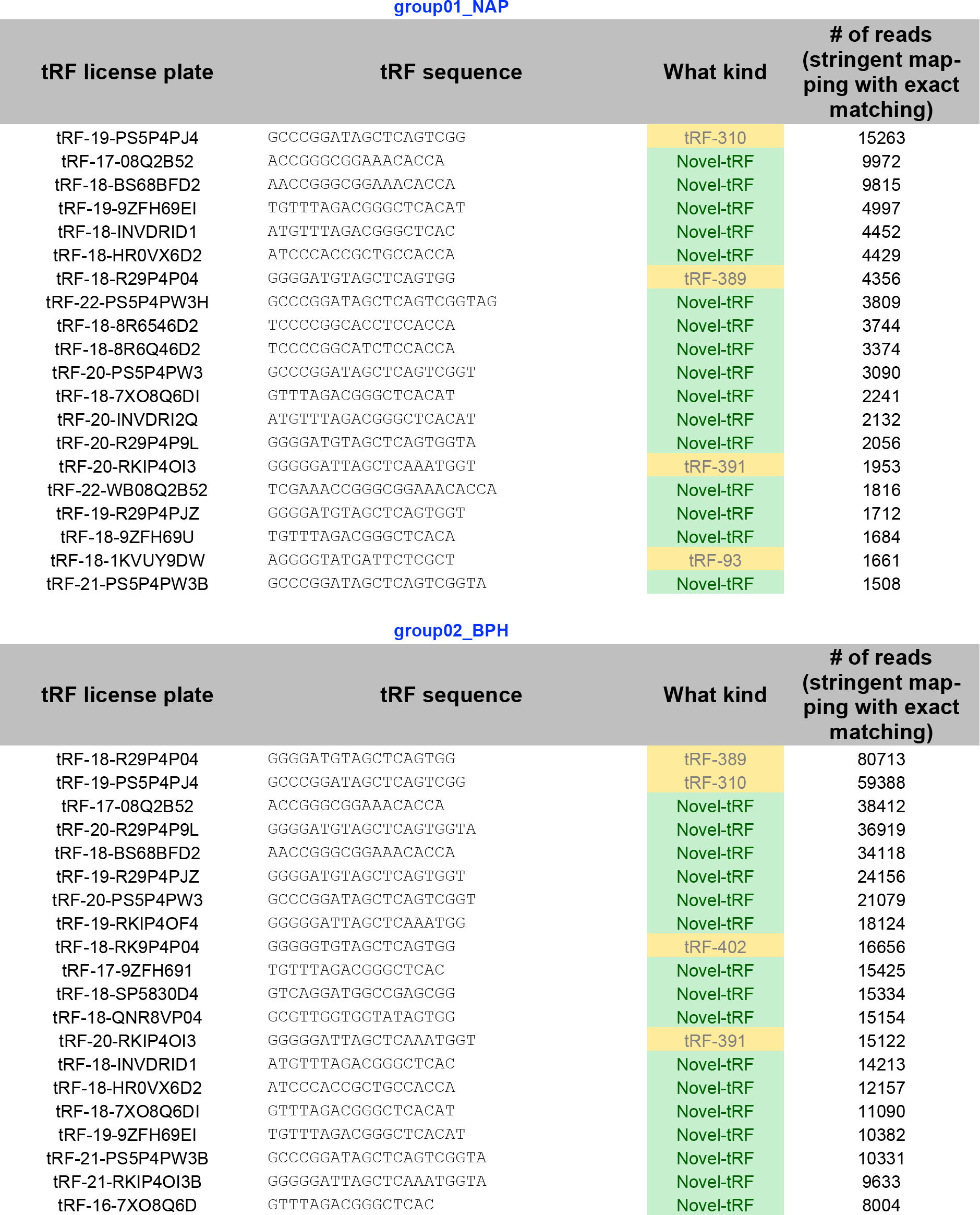
The top-20 most abundant tRFs in each of the 11 datasets of [1] contain many unreported tRFs. For each tRF, the number of reads (stringent, exact mapping) that support it is reported. Also shown are the tRFs’ “license plates” that we described in [6]. The tRFs that were reported in [1] are highlighted in yellow and their label is listed on the third column; otherwise, they are labeled “Novel-tRF”. For the complete list of 3,341 discovered tRFs see Supp. Table 2.

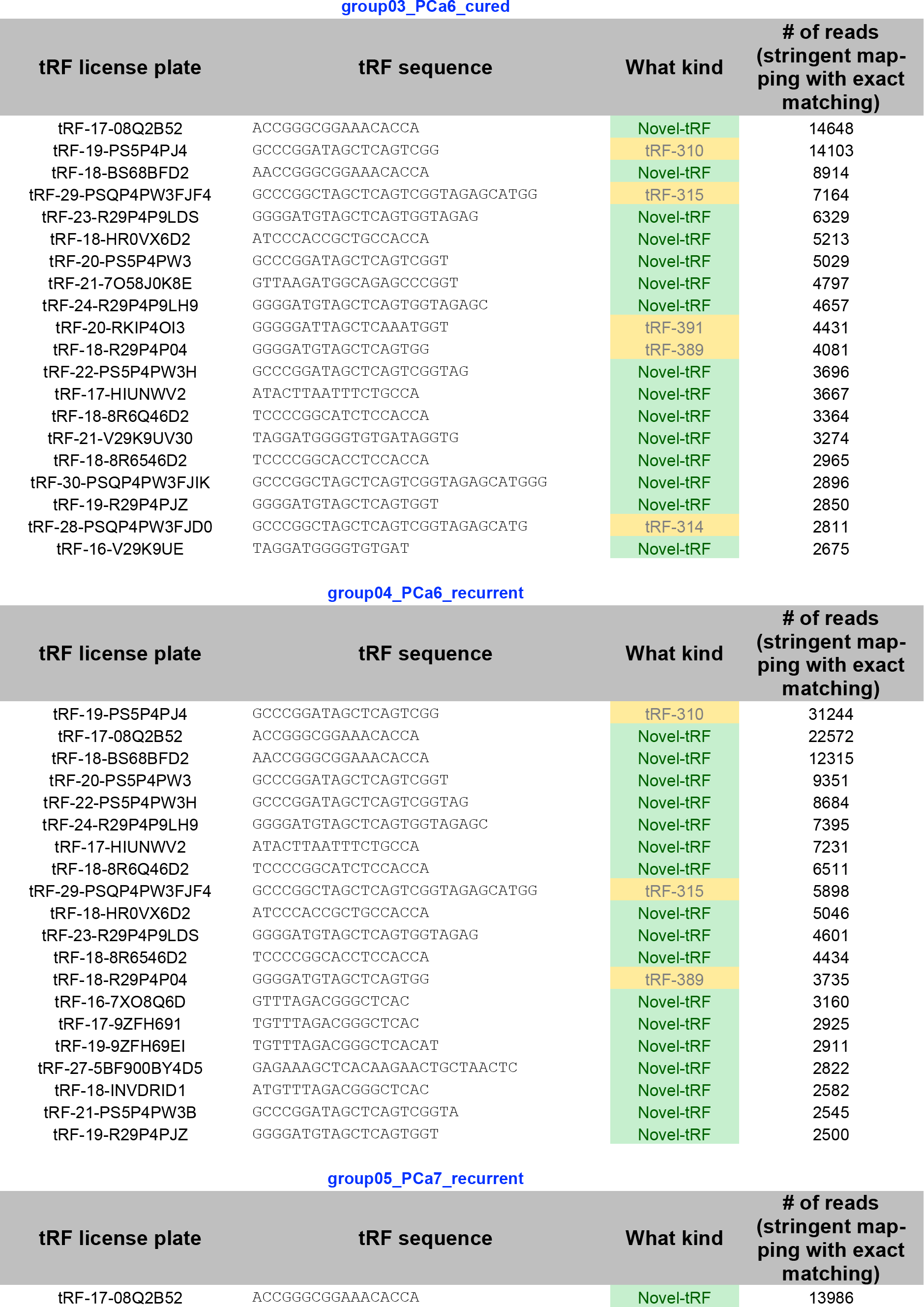

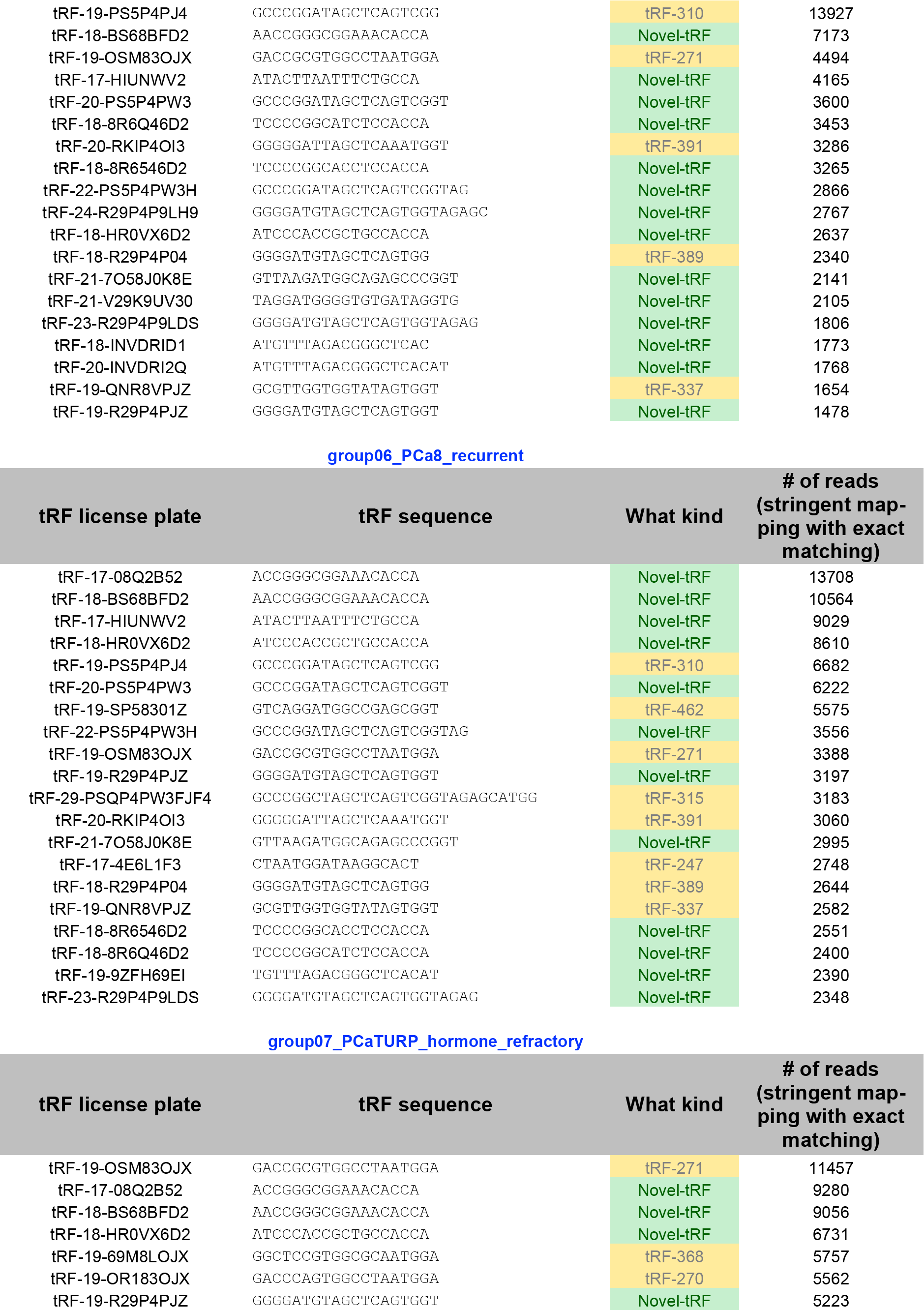

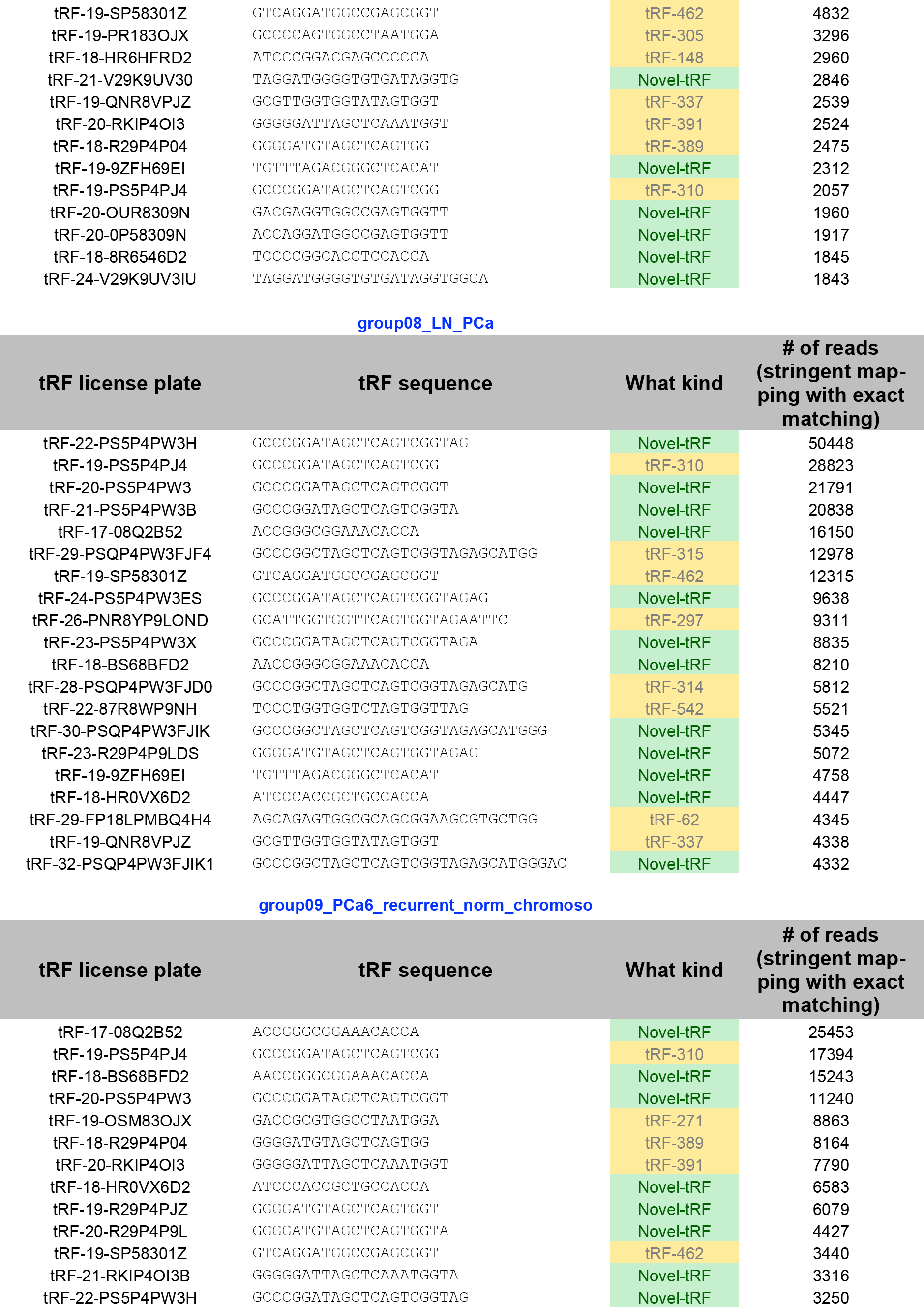

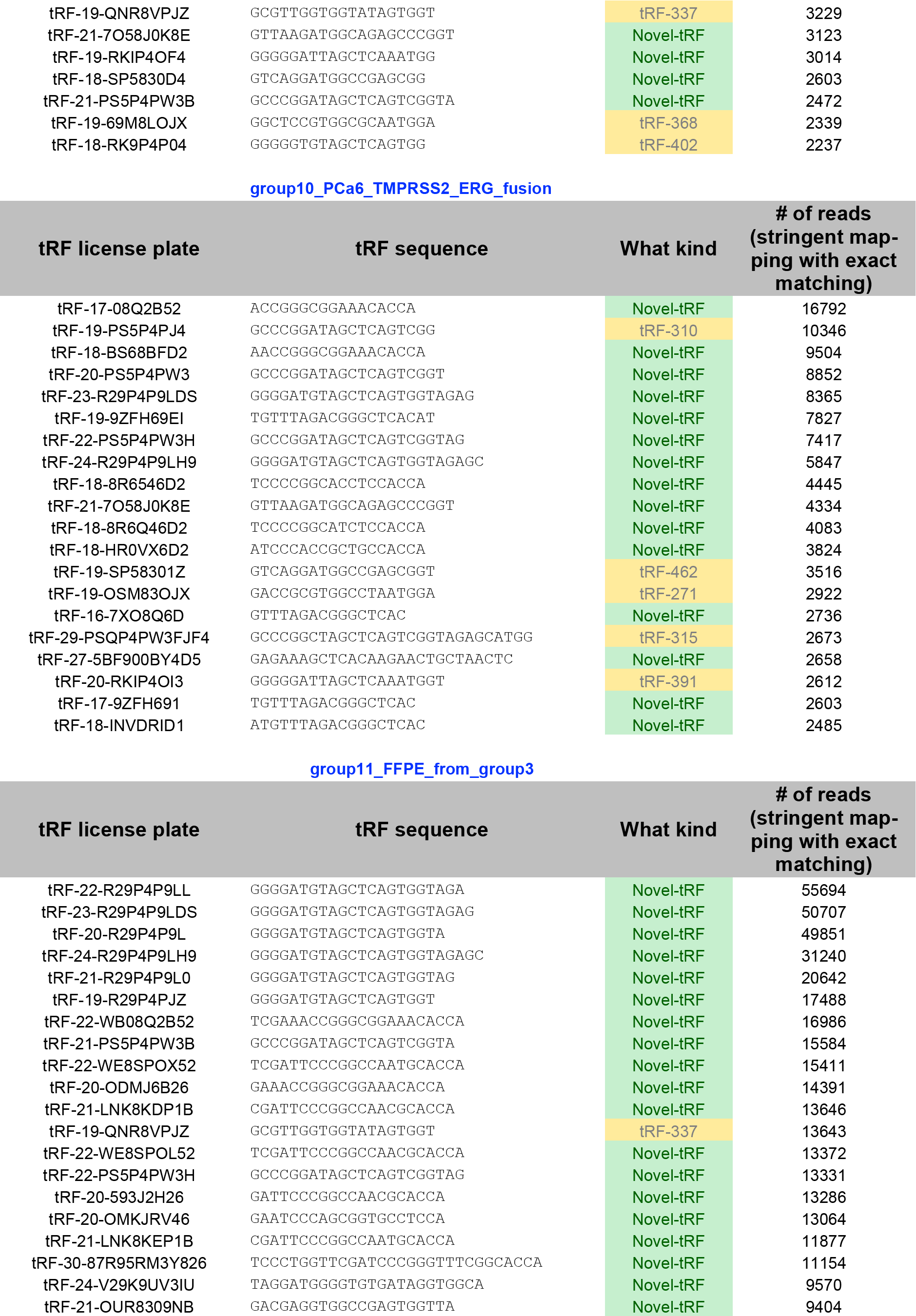

**Table 4.**
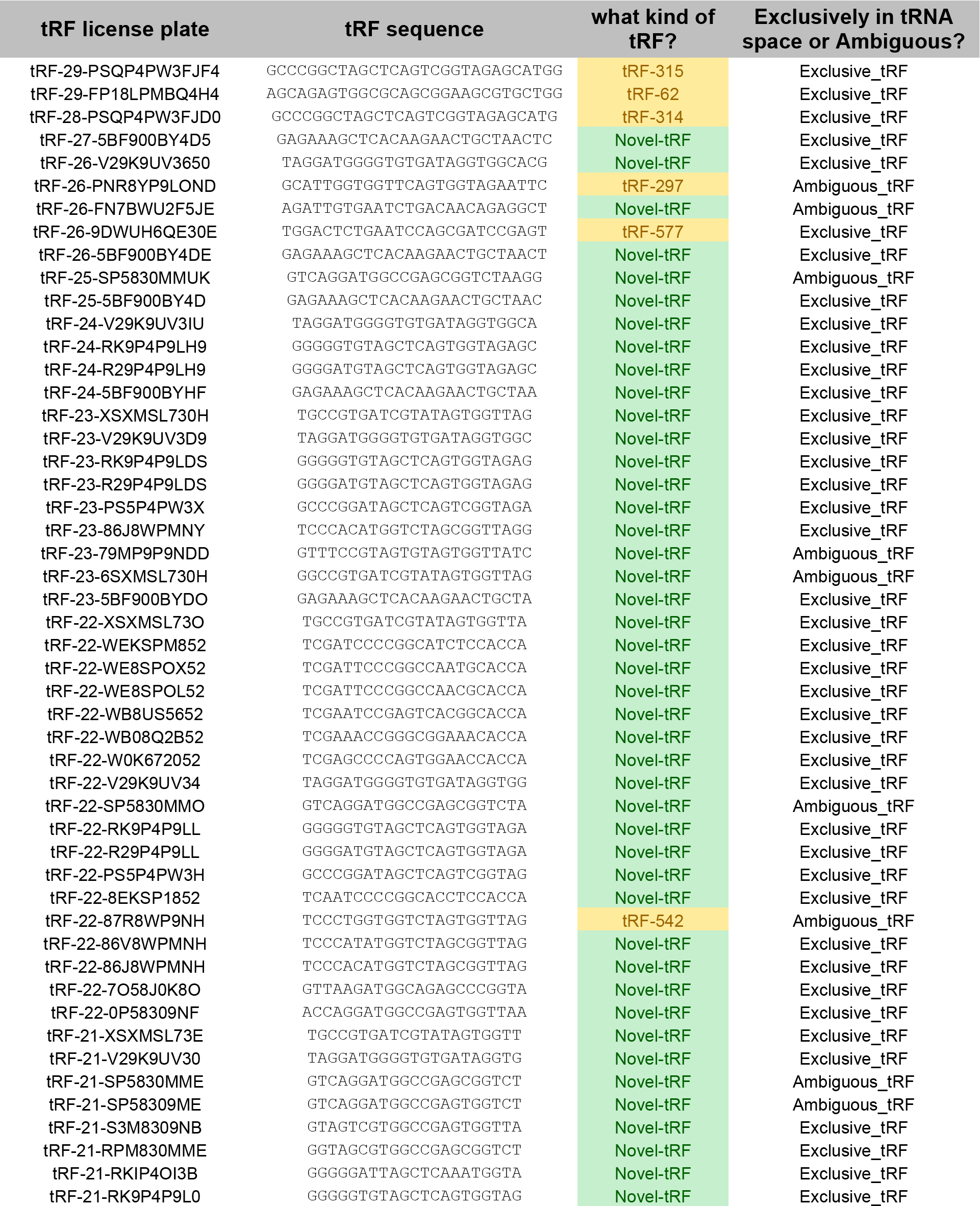
Unreported tRFs that are present in all 11 RNA-seq datasets. Of the 3,341 discovered tRFs, 195 are present in all 11 of the datasets discussed in [1]; however, only 37 of these 195 tRFs were reported in [1]. The below list shows the sequences of all 195 indicating the 37 that were reported. Also shown is whether the tRF is present exclusively in tRNA space or not. For the complete list of 3,341 tRFs and their presence across the 11 RNA-seq datasets of [1] see Supp. Table 3.

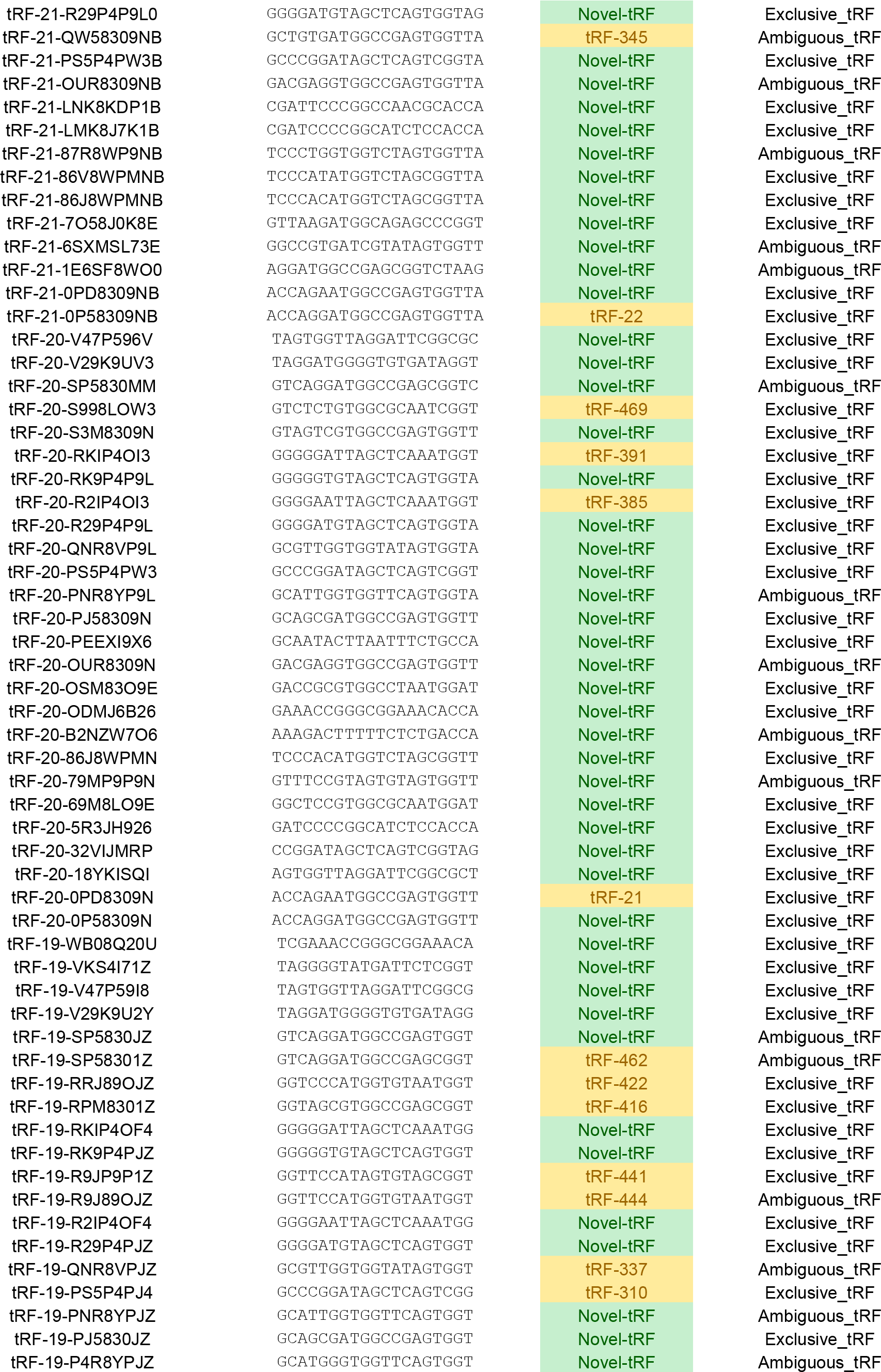

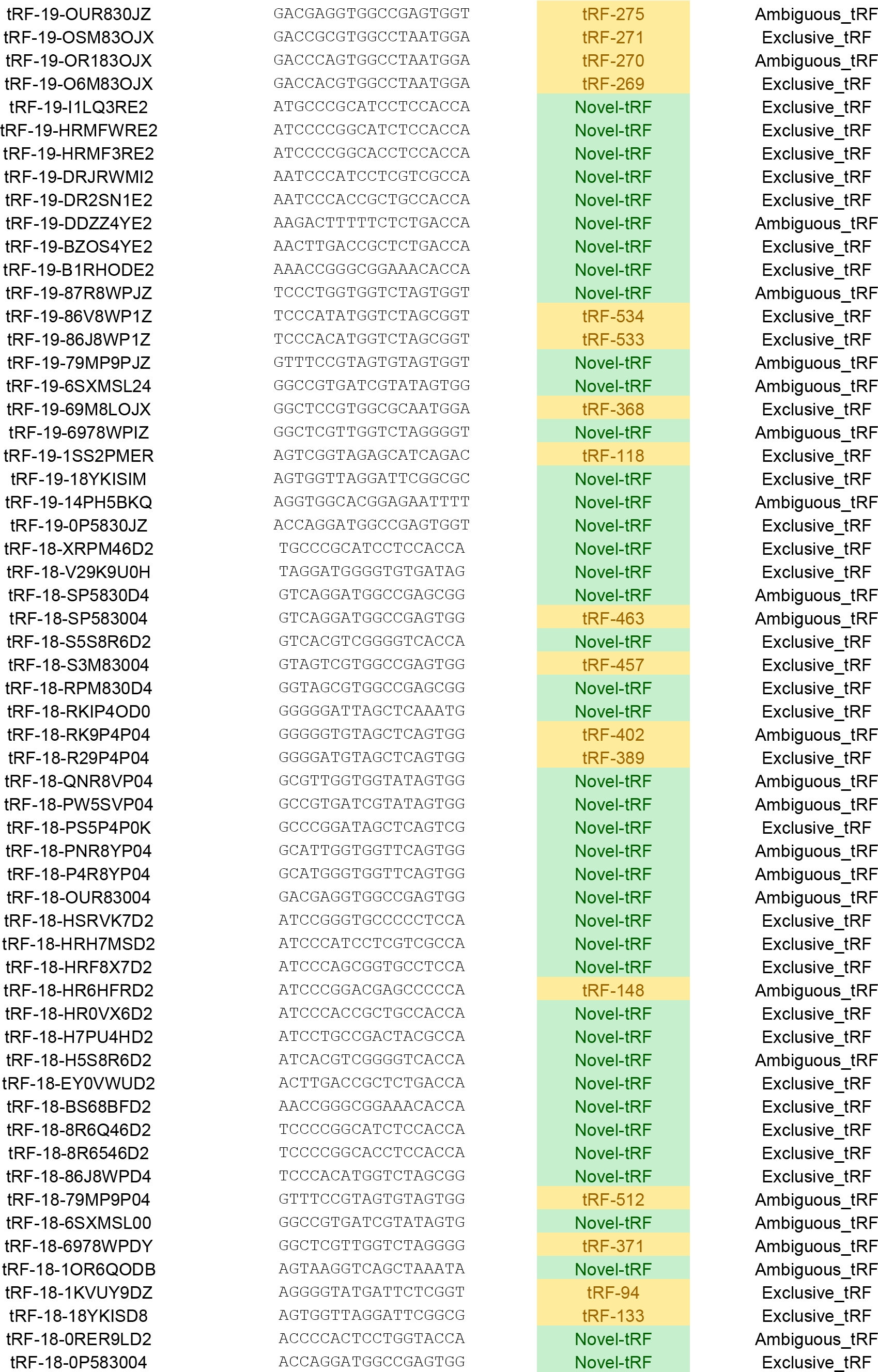

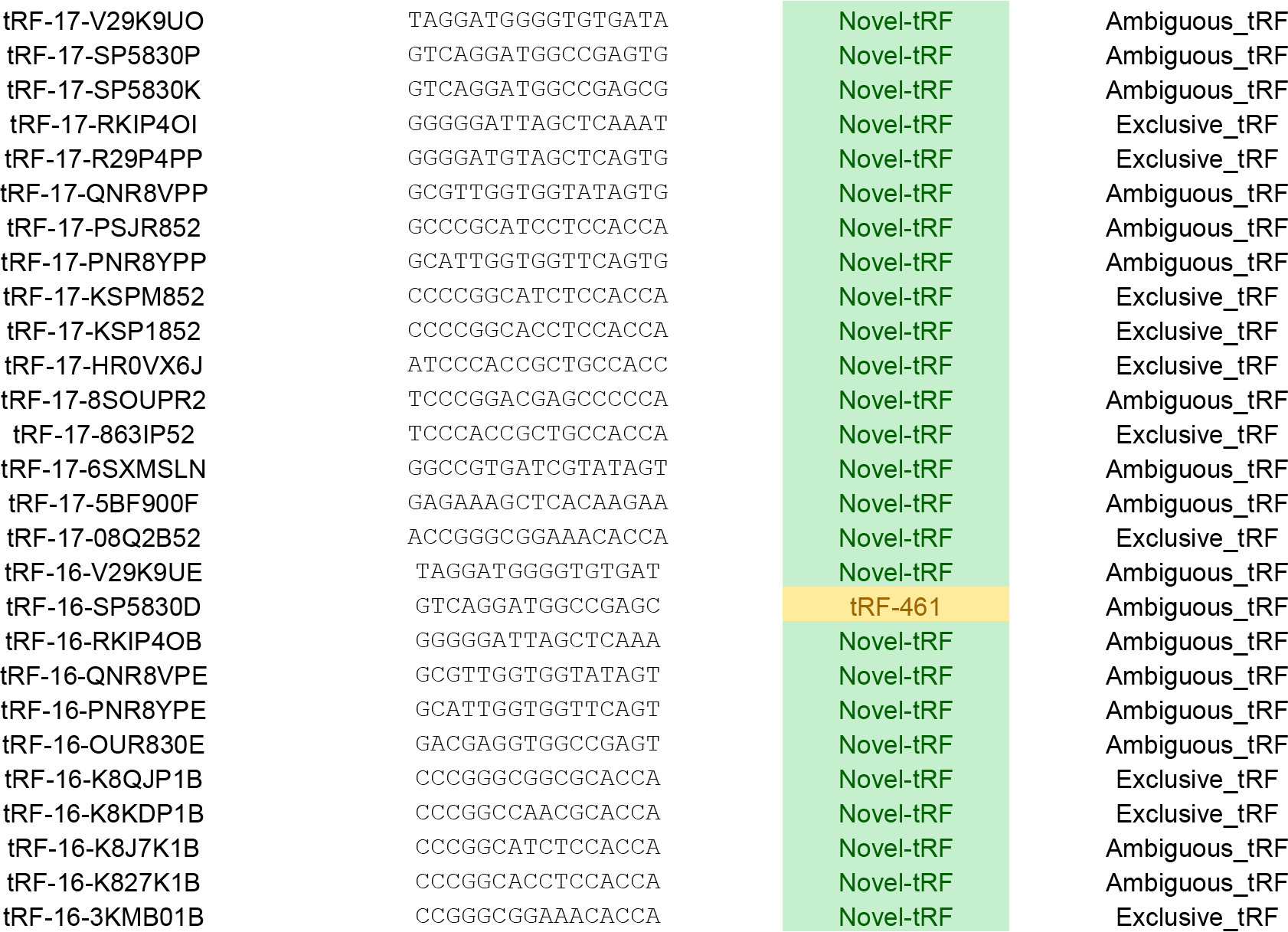

## Many of the tRFs in prostate cancer have sequence relationships that hinder their quantification with conventional q-PCR

Olvedy et al proposed using custom-made locked-nucleic-acid (LNA) primers to quantitate select tRFs. The LNA technology, as practiced in [1], can only guarantee the 3’ termini of target RNAs; thus, it is appropriate for quantifying 5’-tRFs whose 5’ do not require confirmation. However, it cannot guarantee that the correct molecule is quantitated in the case of CCA-ending 3’-tRFs, or, in the case of fully-nested tRFs that have identical suffixes. For example, tRF-185 **CAGTCGGTAGAGCATGGGAC** from [1], which receives 98 reads (stringent mapping) in the LN_PCa dataset, is fully contained in and cannot be distinguished via LNA q-PCR from GCCCGGCTAGCT**CAGTCGGTAGAGCATGGGAC** that is also present in the LN_PCa dataset and receives 4,332 reads (44x more support) – all shown sequences are listed in 5’→3’ orientation. Of the 3,341 tRFs that are identified under stringent discovery conditions, 1,627 (48.7%) form 414 groups each of which contains 2 or more fully-nested tRFs with identical suffixes (Supp. Table 4); for the reasons mentioned above, these tRFs cannot be quantitated using the LNA approach of [1]. Table 5 shows a characteristic example of 20 nested tRFs, from among the 3,341 discovered ones, all of which share the suffix CCGGCTCGAAGGACCA. The complete list of nested tRFs in the profile human prostate cancer data and the corresponding groups is shown in Supp. Table 4.

**Table 5.**
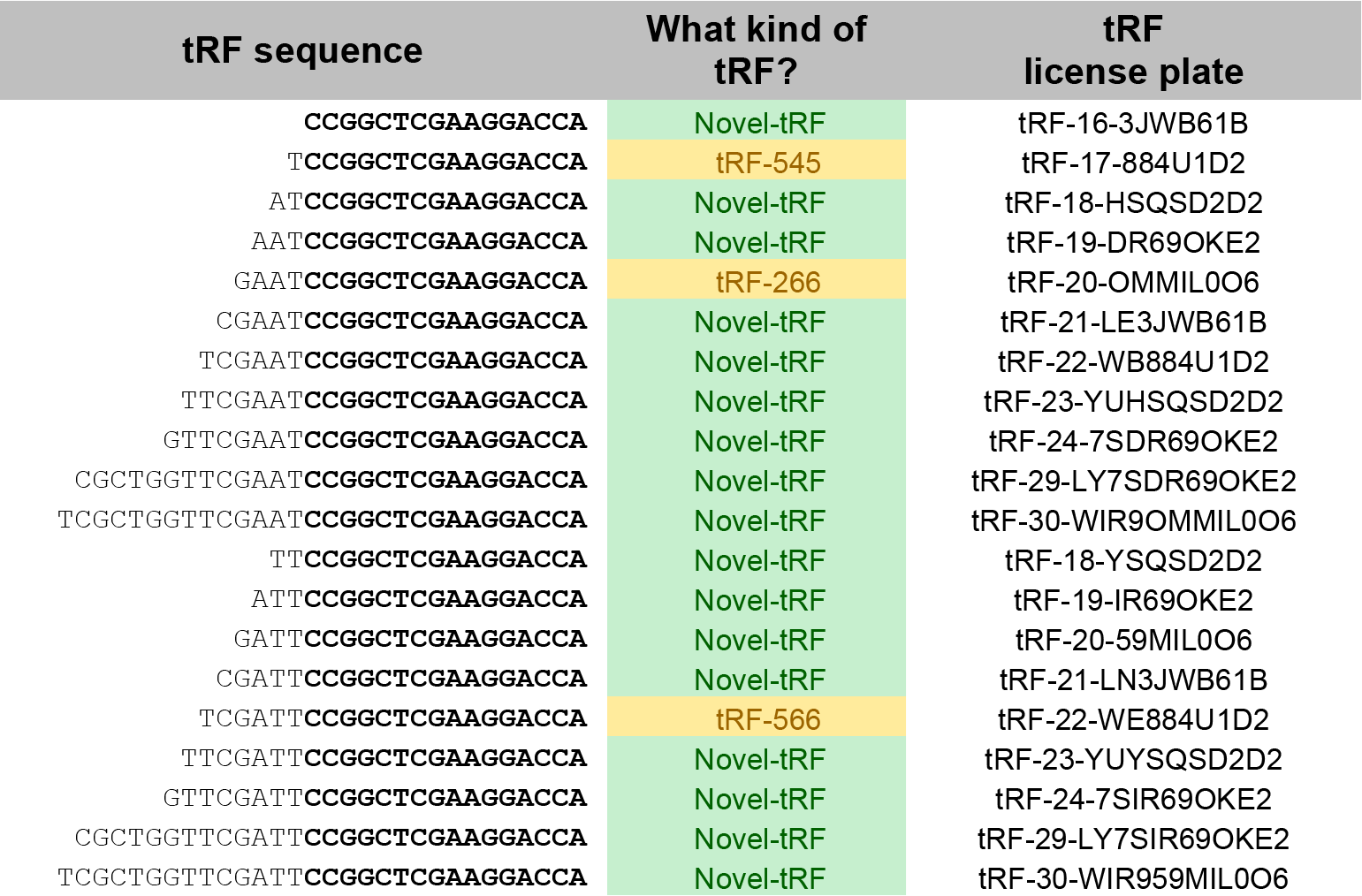
Nested-tRFs in prostate cancer cannot be quantified using the LNA-based method described in [1]. A small example showing nested-tRFs that share a common suffix and are present in the RNA-seq datasets of [1]. The common suffix is shown in boldface. This group of 20 tRFs is one of 414 such groups comprising a grand total of 1,627 of the 3,341 tRFs that are present in the 11 datasets of [1]. The full collection of nested-tRFs is listed in Supp. Table 4.

**NOTE:** the properties of the various tRFs discussed in [1] and of the tRFs that are reported in the Supplemental Tables in the context of the above Discussion can be verified interactively by accessing MINTbase [6] at http://cm.jefferson.edu/MINTbase.

## Authors’ contributions

RM, IR and PL carried out the described analyses. IR, RM, YK, AGT and PL wrote the commentary. All authors read and approved the final manuscript.

## SUPPLEMENTAL MATERIAL

### Supplemental Table l.xlsx

<u>Description:</u> Examples of 63 tRFs from the collection of 598 reported in [1] that have instances in repeat elements that are neither full-length tRNAs nor partial tRNAs.

### Supplemental Table 2.xlsx

<u>Description:</u> List of all the tRFs that are present in the 11 RNA-seq datasets of [1]. The discovered tRFs are reported separately for each dataset.

### Supplemental Table 3.xlsx

<u>Description:</u> For each of the 3,341 tRFs we discovered in the 11 RNA-seq datasets of [1], the Table shows the number of different datasets in which each tRF is present.

### Supplemental Table 4.xlsx

<u>Description:</u> A list of tRFs that cannot be quantitated with the LNA-based approach of [1]. The Table lists 414 groups comprising 1,627 of the 3,341 tRFs that were discovered in the 11 RNA-seq datasets of [1]. Within each group, tRFs are fully-nested in various combinations while sharing identical suffixes.

## REFERENCES

1. Olvedy M., et al., A comprehensive repertoire of tRNA-derived fragments in prostate cancer. Oncotarget, 2016 - [Epub ahead of print].

2. Keam, S.P. and G. Hutvagner, tRNA-Derived Fragments (tRFs): Emerging New Roles for an Ancient RNA in the Regulation of Gene Expression. Life (Basel), 2015. 5(4): p. 1638–51.

3. Raina, M. and M. Ibba, tRNAs as regulators of biological processes. Front Genet, 2014. 5(298): p. 171.

4. Honda S., et al., Sex hormone-dependent tRNA halves enhance cell proliferation in breast and prostate cancers. Proc Natl Acad Sci U S A, 2015. 112(29): p. E3816–25.

5. Telonis, A.G., et al., Dissecting tRNA-derived fragment complexities using personalized transcriptomes reveals novel fragment classes and unexpected dependencies. Oncotarget, 2015. 6(28): p. 24797–822.

6. Pliatsika V., et al., MINTbase: a framework for the interactive exploration of mitochondrial and nuclear tRNA fragments. Bioinformatics, 2016 - [Epub ahead of print].

7. Telonis, A.G., et al., Consequential considerations when mapping tRNA fragments. BMC Bioinformatics, 2016. 17(1): p. 123.

